# Deciphering sex-specific miRNAs as heat-recorders in zebrafish

**DOI:** 10.1101/2021.12.29.474410

**Authors:** T.A. van Gelderen, J. Montfort, J.A. Álvarez-Dios, V. Thermes, F. Piferrer, J. Bobe, L. Ribas

## Abstract

MicroRNAs (miRNAs) are important post-transcriptional regulators of gene expression in a wide variety of physiological processes, including those related to the reproductive system. Although in the last decade a plethora of miRNAs has been reported, the miRNA alterations occurred by environmental cues and their biological functions have not yet been elucidated. With the aim to identify epigenetic regulations mediated by miRNAs in the gonads in a climate change scenario, zebrafish (*Danio rerio*) were subjected to high temperatures during sex differentiation (18-32 days post fertilization, dpf), a treatment that results in male-skewed sex ratios. Once the fish reached adulthood (90 dpf), ovaries and testes were sequenced by high-throughput technologies. About 101 million high-quality reads were obtained from gonadal samples. Analyses of the expression levels of the miRNAs identified a total of 23 and 1 differentially expressed (DE) miRNAs in ovaries and testes, respectively, two months after the heat treatment. Most of the identified miRNAs were involved in human sex-related cancer. After retrieving 3’ UTR regions, ~400 predicted targets of the 24 DE miRNAs were obtained, some with reproduction-related functions. Their synteny in the zebrafish genome was, for more than half of them, in the chromosomes 7, 2, 4, 3 and 11 in the ovaries, chromosome 4 being the place where the predicted sex-associated-region (*sar*) is localized in wild zebrafish. Further, spatial localization in the gonads of two selected miRNAs (miR-122-5p and miR-146-5p) showed exclusive expression in the ovarian germ cells. The present study expands the catalog of sex-specific miRNAs and deciphers, for the first time, thermosensitive miRNAs in the zebrafish gonads that might be used as potential epimarkers to predict environmental past events.

## Introduction

Water sea temperature levels have been rising in the last 60 years [1], with critical consequences for marine and aquatic life. Fish, thanks to their thermal plasticity, are able to survive the variations of water temperatures [2]. Nevertheless, sex determination in fish, unlike in mammals, is regulated by genetic and environmental factors [3,4], and consequently, higher temperatures during sex differentiation skew the sex ratio towards males in many fish species [5]. Since the first study showing the crosslink between masculinization occurred by heat treatments and DNA methylation in the European sea bass (*Dicentrarchus labrax*) gonads [6], in the last decade, studies describing the role of epigenetics in sexual development have emerged. In pufferfish (*Takifugu rubripes*), DNA methylation alterations were able to faithfully describe dimorphic differences in the gonadal epigenomes of fish subjected to different thermal regimes, identifying two genes (*amhr2* gene and *pfcyp19*) as main actors of sex determination in this species [7]. Similarly, in half-smooth tongue sole (*Cynoglossus semilaevis*) differentially methylated regions (DMR) were observed in the gonads of sex-reversed fish indicating that high-temperature treatments override sexual fate determined by genetic factors through epigenetic pathways [8]. Recent transgenerational studies in zebrafish (*Danio rerio*) showed that temperature affected the testicular epigenome in the first generation but these effects were washed out in the second generation [9].

miRNAs are small, non-coding RNAs, consisting of approximately 22 nucleotides, and are considered as epigenetic mechanisms responsible to regulate the post-transcriptional cellular machinery. These molecules regulate gene expression by preventing protein translation through binding to their target messenger RNAs (mRNA), serving as recruiters in the mRNA degradation pathways [10]. Over the past two decades, miRNA-related research has expanded considerably, as in only two years the miRNA submissions in public databases increased by fifty percent [11]. Over 3,500 mature miRNAs have been identified in 16 teleostei species (www.mirbase.org), zebrafish being the first fish species with more miRNAs described in detail [12]. In fish, miRNAs play pleiotropic functions, for example, in the reproduction system, immune system, metabolism, and skeletal formation, among others (reviewed in [13]). In the last few years, studies in adult fish have revealed the presence of sexual dimorphism in the miRNA expression between ovary and testis in some species such as [14], yellow catfish (*Pelteobagrus fulvidraco*) [15], rainbow trout (*Oncorhynchus mykiss*) [16], tilapia (*Oreochromis niloticus*) and zebrafish [17,18].

Since miRNA alterations can respond to environmental influences, they are foreseen as potential targets for improving productivity in aquaculture. Some studies have addressed the temperature effects on different target tissues. miRNA expression changed in response to increasing natural temperatures in zebrafish embryonic fibroblast cells [19] and in rainbow trout liver [20] and head kidney [21]. Cold tolerance has been tested by detailing miRNA expression in Emerald rockcod (*Trematomus bernacchii*) gills [22], in turbot (*Scophthalmus maximus*) brain, head kidney and liver [23], and in sole (*Solea senegalensis*) embryos [24]. To date, only few studies have addressed the miRNA alterations due to temperature increases in the gonads. Juvenile Atlantic cod showed some, but few, DE miRNAs after heat during early development, although no differences between ovaries and testes were addressed [25]. Further, adult zebrafish gonads subjected to high temperature in combination with antidepressant compounds showed a variation of the miRNA abundance by a target miRNA approach [26]. Thus, to our knowledge, no data regarding the long-term effects of the high temperatures in the miRNome of the ovaries and testes in fish have ever been reported. Therefore, the goal of this study was to characterize a set of miRNAs that could be used as epimarkers of the effects of heat-stress on fish gonads in a context of global warming.

## Materials and methods

### Experimental design

The AB zebrafish were reared at the experimental aquarium facilities of the Institute of Marine Sciences (ICM-CSIC) in Barcelona. Fish husbandry and thermal treatments were done as previously described in Ribas *et al*. [27]. For this experiment, ~175 spawned eggs by a single pair mating were used. At 6 days post fertilization (dpf), 35 larvae were equally distributed into four tanks (two technical replicates for each group) of 2.8 liters (Aquaneering, mod. ZT280) to avoid high-density masculinization effects [28]. Fish were exposed to high temperature (HT) at 34 ± 0.5°C or to control temperatures (CT) at 28 ± 0.5°C between 18-32 dpf. The temperature was changed at a rate of 1.5°C/day to reach the desired temperatures. After the heat treatment, animals were grown until gonadal maturation, i.e., 90 dpf. The Chi-squared test with arcsine transformation was used to study differences in sex ratios. Biometry differences between CT and HT were determined by Student *t*-tests. Previously, for each group, homoscedasticity of variances and normality were checked by Levene’s test and Shapiro-Wilk test, respectively.

### Sampling, sample selection and RNA extractions

Adult fish were sacrificed by cold thermal shock and the sex of the fish was visually assessed under the microscope. Gonads were isolated and flash-frozen into liquid nitrogen and kept at −80°C for further analyses. To unify the gonadal maturation, samples were selected based on two criteria. First, based on macroscopical examination following Ribas *et al*. [27], and second, based on the highest gene expression levels of gonadal aromatase (*cyp19a1a*) and anti-Müllerian hormone (*amh*) in ovaries and testes, respectively, that worked as sex-markers (data not shown) [29,30]. miRNA of sixteen gonads (four samples each sex and treatment) was isolated by miRNAs isolation commercial kit (Qiagen^®^ miRNA, 217004) and quality was assessed by BioAnalyzer (2100 Bioanalyzer, Agilent Technologies). On average, RNA Integrative Number (RIN) values for all the samples were ≥ 9, indicating high score RNA qualities.

### Small RNA library and sequencing

In total, 16 libraries were constructed individually from zebrafish gonads. Library preparation was performed by NEBNext^®^ Small RNA Library Prep Set for Illumina^®^ (Multiplex Compatible) kit following manufacturer’s instructions. Sequencing (1×50, v4, HiSeq) was performed at single-end mode with a read length of 50 bp at the Genomics Unit of the Centre for Genomic Regulation (CRG) in Barcelona.

### miRNA validation and gene expression analyses

Validation of the miRNA sequencing data was done by RT-qPCR of those selected sequenced miRNAs. cDNA was generated using the miRNA 1st-Strand cDNA Synthesis Kit (Agilent Technologies) following manufacturer’s instructions. Firstly, the polyadenylation reaction was performed after cDNA synthesis. qPCR was performed using the qPCRBio SyGreen blue mix low ROX (PCR Biosystems). A mix of 5 μL 2x qPCRBIO SyGreen Blue mix, 0.4 μL forward primer, 0.4 μL universal reverse primer (Agilent Technologies), 100 ng cDNA and H_2_O up to 10 μL was made for each sample. The sequences of the forward primers for the six selected miRNAs were as follows: dre-miR-202-5p: TTCCTATGCATATACCTCTTT, dre-miR-92a-3p: TATTGCACTTGTCCCGGCCTGT, dre-miR-21-5p: TAGCTTATCAGACTGGTGTTGGC, dre-miR-146b-5p: TGAGAACTGAATTCCAAGGGTG, dre-miR122-5p: TGGAGTGTGACAATGGTGTTTG, dre-miR-2189-3p: TGATTGTTTGTATCAGCTGTGT. The dre-U6 (ACTAAAATTGGAACGATACAGAGA) was used as the reference gene. The comparisons for validations were performed as follows: ovary high temperature (OHT) *vs*. ovary control temperature (OCT) for dre-miR-202-5p, dre-miR-92a-3p, dre-miR-21 and dre-miR-146b-5p; testis high temperature (THT) *vs*. testis control temperature (TCT) for dre-miR-122-5p; OCT *vs*. TCT for dre-miR-146b-5p and dre-miR-2189-3p.

### Bioinformatics: miRNA mapping and annotations

Sequenced libraries were analyzed by Prost! as described by Desvignes *et al*. [18]. Briefly, sequencing data were trimmed and reads were mapped on the reference genome version 11 of zebrafish (GRCz11) for annotations and to distinguish novel and known miRNAs. Expression of these miRNAs was determined by the raw count matrix used as input into DESeq2. Read data were normalized by DESeq functions and relative expression between groups was generated by base mean, log2 fold change and adjusted p-value (*P* < 0.05). To visualize the level of similarity of individual samples a Multi-Dimensional Scaling (MDS) plot was created with the package EdgeR [31,32] from Bioconductor [33] Heatmaps of DE miRNAs (HT *vs*. CT and O *vs*. T) were constructed using the R package pheatmap (https://CRAN.R-project.org/package=pheatmap).

### Consistent gonadal miRNAs in zebrafish

To identify miRNAs in the zebrafish gonads, miRNA data from two available publications in zebrafish was used [17,18], additionally to the data currently presented. The normalized read lists were used to identify significantly expressed miRNAs in testes or ovaries with an expression of 100 normalized reads or higher. Next, DE miRNAs between ovaries and testes in the three miRNA datasets were identified (expression of 100 normalized reads or higher and adjusted *P* ≤ 0.05). Venn Diagrams were created using the software from the Bioinformatics & Evolutionary Genomics group from Ghent University (http://bioinformatics.psb.ugent.be/webtools/Venn/).

### Functional annotation of miRNA targets

To identify miRNA targets, 3’UTR regions and genome annotation for zebrafish were extracted from Ensembl (https://www.ensembl.org/) using the Biomart data mining tool. Putative miRNA targets were identified with MiRanda [34] with energy threshold −25 and other parameters left to their default value. Subsequent MiRanda output was pruned and processed to extract relevant information with a custom Perl script. The portal (https://david.ncifcrf.gov/) was used to perform enrichment analyses and search for GO terms and KEGG pathways. Graphs of a representative summary of each GO term category for each gonad class were produced with Revigo [35] using term frequency as the guiding parameter. A circular zebrafish genome graph was produced with Circos [36]. MDS samples graph was created with package edgeR [31] from Bioconductor [33]. When necessary, custom Perl scripts were created to extract information and combine data throughout the bioinformatic analyses.

### Fluorescent in situ hybridization

For fluorescent *in situ* hybridization (FISH), ovaries dissected from a total of 12 zebrafish adult females were fixed overnight in 4% paraformaldehyde (PFA) at 4°C, dehydrated in 100% methanol and stored at −20°C. Fixed ovaries from the control group (OCT) were paraffin-embedded and sections (9 μm thickness) were obtained with a microtome (HM355, microm). The anti-sense miRCURY LNA miRNA detection probe dre-miR-146b-5p (YD00613622, QIAGEN) was used. The mmu-miR-122-5p miRCURY LNA probe (YD00615338, QIAGEN) was used to detect the dre-miR-122-5p mature form, since zebrafish and mouse miR-122-5p sequences are identical. Probe sequences were 5’-CAAACACCATTGTCACACTCC-3’ and 5’-CACCCTTGGAATTCAGTTCTC-3’ to detect dre-miR-122-5p and dre-miR-146b-5p, respectively. The scramble-miR miRCURY LNA Detection probe (5’-GTGTAACACGTCTATACGCCCA-3’, YD00699004, QIAGEN) was used as a negative control. All LNA probes were double-DIG labeled at both 5’ and 3’ ends. FISH was performed using the miRCURY LNA miRNA ISH kit (FFPE, 339450, QIAGEN) following the manufacturer’s instructions, Permeabilization was performed for 7 min at room temperature using Proteinase-K (10 μg/ml, P2308 Sigma). LNA probes were used at 40 nM at 53°C (30°C below the RNA Tm) for 2 h. Samples were then incubated overnight at 4°C with a rabbit anti-DIG HRP-conjugate antibody (1:500, Roche). Then, the anti-DIG-HRP antibody was detected with the TSA-Cy3 substrate (1:50, TSATM PLUS Cy3 kit, NEL 745001KT, Perkin Elmer) for 10 min at room temperature. Nuclei were stained with 4% Methyl Green (MG, 323829-5G, Sigma-Aldrich) in PBS/0.1% triton for 15 min at room temperature All pictures were taken with a Leica TCS SP8 laser scanning confocal microscope using 552 nm and 638 lasers for TSA-cy3 and MG detection, respectively.

## Results

### Sex ratio and biometry

After heat treatment during sex differentiation, a 17% masculinization was observed in the high temperature (HT) group (S1 Fig), although differences were not significant. The accumulated degrees during the treatment were 419.84 and 511.25 for control temperature (CT) and HT, respectively. The mean weight of the animals was as follows: female CT 0.38± 0.09g, male CT 0.27± 0.08g, female HT 0.26 ± 0.10g and male HT 0.24± 0.09g. The mean size of the animals was as follows: female CT 2.48±0.26 cm, male CT 2.48±0.05 cm, female HT 2.53±0.15 cm, male HT 2.35±0.10 cm. No significance was found in either female or male CT *vs*. HT in weight or size. The weights, lengths and K-factor and statistical results of all 16 fish can be found in S1 Table.

### miRNA sequencing overview and validation

On average, we obtained 25.3 million sequences per library and the total number of sequences exceeded 101 million, 71 and 30 million for testes and ovaries, respectively. The length distribution showed that over 99% of the obtained sequences were ~30 nucleotides (nts), with a range between 36–37 nts in length. A total of 359 mature miRNAs were identified after alignments against the zebrafish genome (Dataset 1). Eleven miRNAs were not fully annotated against the zebrafish genome, of which four were aligned to other fish species (miR-122-3p in 20 species, one miRNA annotated as let-7a/c/e/f/k-3p in 18 species, one miRNA annotated as let-7e/f/g-2-3p in 10 species and another one as miR-139-5p in 16 species). Only seven miRNAs were not annotated and were defined as novel. Thus, 98.1% of the miRNA sequenced was annotated. The raw sequencing data were made publicly available in NCBI SRA with the accession number: PRJNA755482.

The MDS analyses clustered the samples based on their corresponding group by sex and treatment (S2 Fig). The two MDS components explained 57% of the variance among the samples. There was one testicular control sample (i.e., TCT3) that was clustered with the heat treated samples, but was not discarded from further analyses. Similarly, one ovarian control sample (OCT5) was grouped among treated ovarian samples and also kept for analysis.

miRNA-seq data was validated by testing the expression of six miRNAs (dre-miR-202-5p, dre-miR-92a-3p, dre-miR-21-5p, dre-miR-146b-5p, dre-miR-122-5p, dre-miR-2189-3p) by qPCR analyses in the ovary and testis in three different comparisons based on their expression in sequencing data. Results showed a linear regression with R^2^ = 0.9522 and *P* = 0.00087, thus validating miRNA sequencing results (S3 Fig).

### miRNAs in the zebrafish gonads

Data from two similar studies from the same zebrafish strain [18] and from a different zebrafish line (crossing nacre transparent, − /−, with zf45Tg [17] were used in order to identify miRNAs that were consistently expressed in the zebrafish gonads within one given sex.

Comparing our miRNA data with the two available libraries, we found 32 and 50 miRNAs in ovary and testis, respectively, specific for our data. A total of 131 and 137 miRNAs in the ovary and the testis, respectively, were found between the results reported by Desvignes *et al* 2019 and our present data while only 37 and 34 were common between our data and the results of Presslauer *et al* 2017 (Fig 1A, B). Between all three libraries, 35 and 32 common miRNAs in ovaries and testes were found, respectively (Fig 1A, B and S2 Table). A total of 25, 14 and 20 DE miRNAs in ovary and 16, 3 and 26 in testis for Presslauer *et al*., Desvignes *et al*. and our data, respectively, were identified as unique for each of the three publications (S4 A, B Fig). Comparing DE miRNAs between ovary and testis in the three studied data, identified 1 common miRNA for each studied tissue (S4A, B Fig), dre-miR-200b-3p in ovary and dre-miR-212-5p in testis. Since our and Desvignes *et al*. 2019 data used the same zebrafish AB strain, we selected those common DE miRNAs between ovary and testis (8 and 11, respectively) to plot a heatmap (Fig 1C) that showed those miRNAs that were constitutively differentially expressed between both sexes.

**Fig 1.**
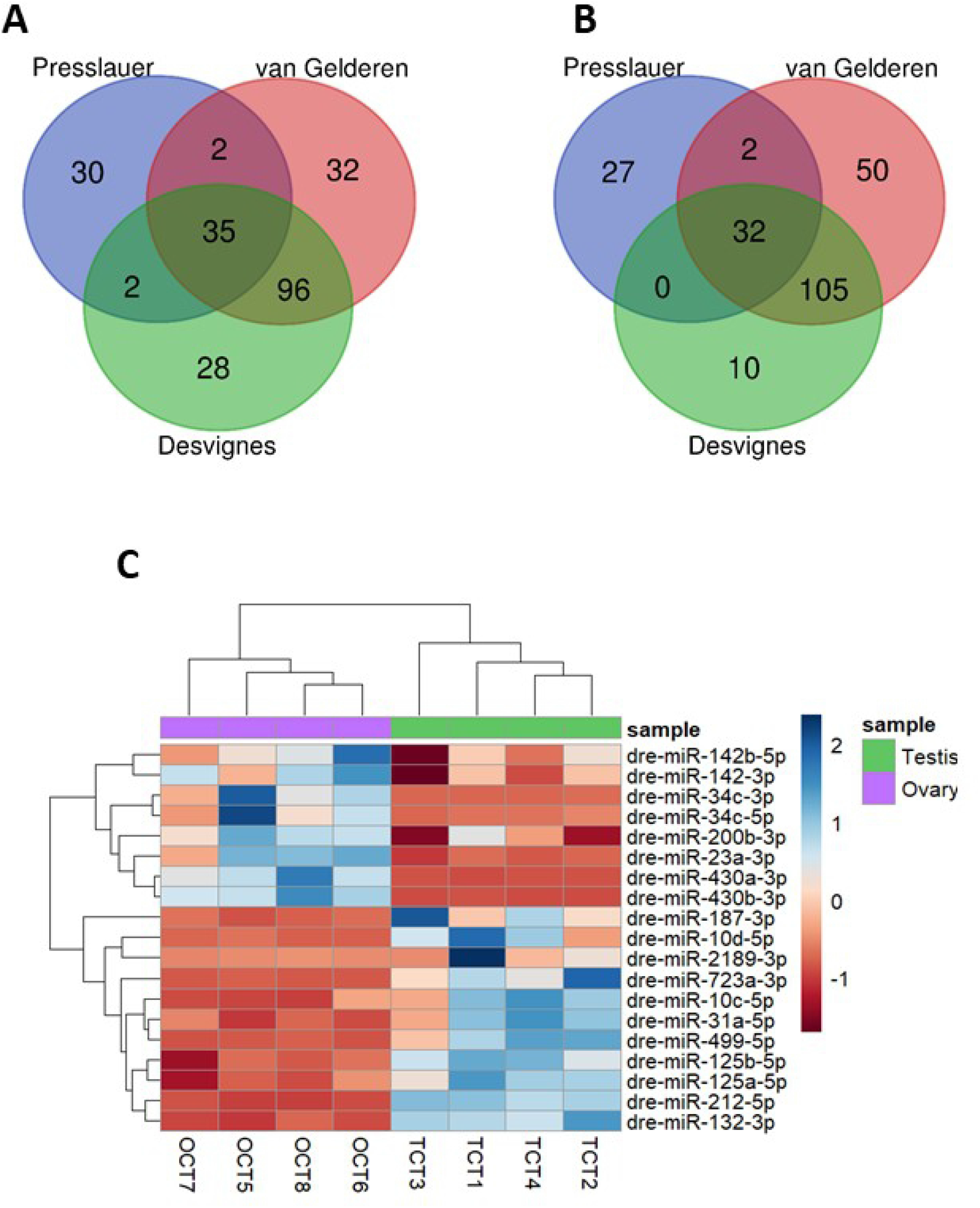
The number of miRNAs that were differentially expressed (DE) of two AB strains (current data and Desvignes *et al* 2019) and zf45Tg hybrid zebrafish (Presslauer *et al* 2019). **A)** In ovaries, and **B)** testes. **C)** Heatmap of DE miRNAs between ovary and testis commonly found in Desvignes *et al* 2019 and present data.

### miRNAs sensitive to temperature in the gonads

One miRNA was found to be significantly upregulated (adjusted P-value ≤0.05) in testis between CT and HT groups, i.e., dre-miR-122-5p, and 23 miRNAs were differentially regulated in the ovary (adjusted *P*-value ≤0.05, Fig 2) (S3 Table), giving a total of 24 DE miRNAs. The five top upregulated miRNAs in the ovary were dre-miR-499-5p, dre-miR-202-5p, dre-miR-92b-3p, dre-miR-454b-3p, and dre-miR-725b-5p. The most downregulated were dre-miR-726-5p, dre-miR-184-3p, dre-miR-146b-5p, dre-miR-34a-5p and dre-miR-132-3p (S5 Fig). The temperature-induced higher fold changes in those downregulated miRNAs compared to those upregulated, show a difference in expression over six-fold.

**Fig 2.**
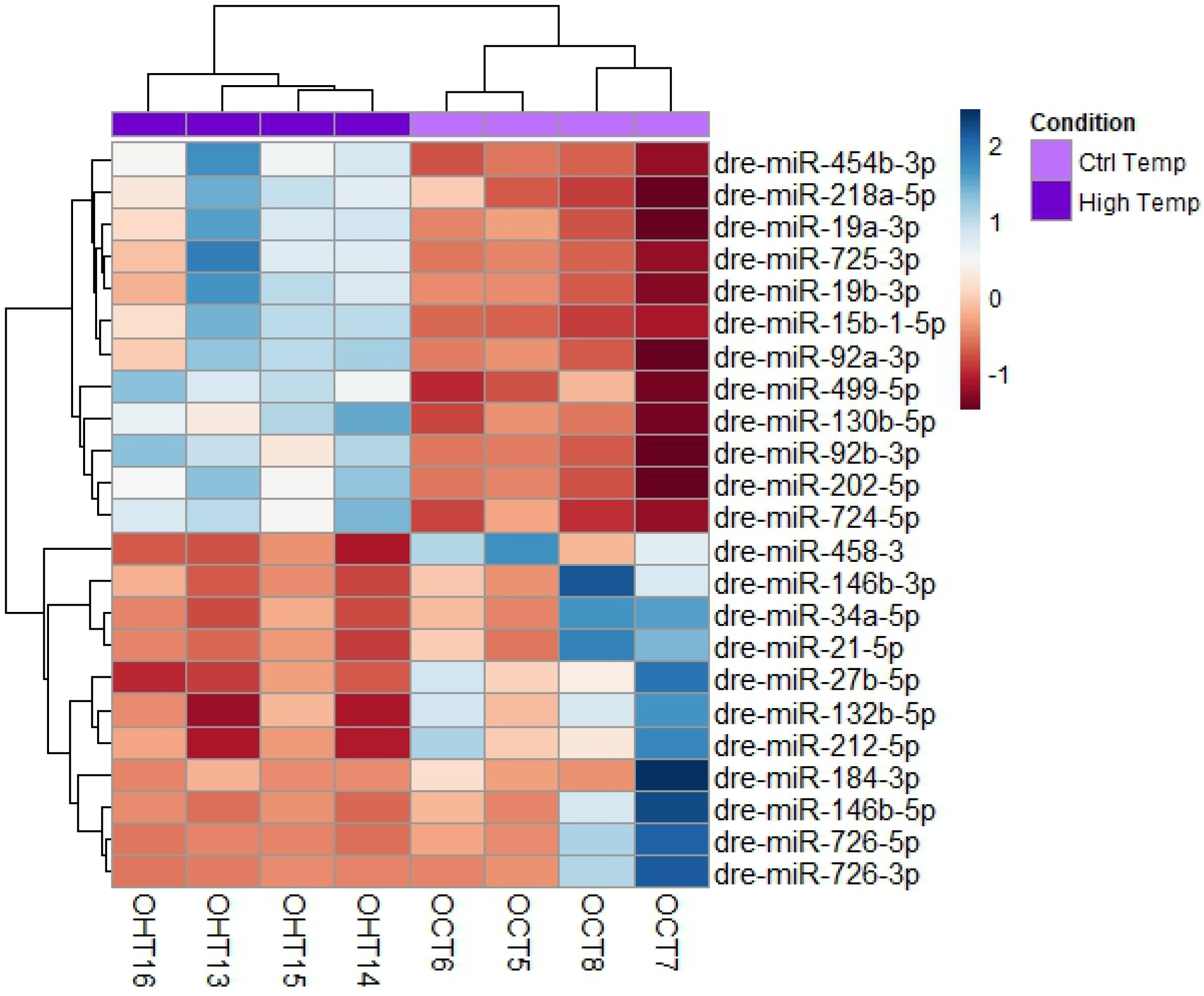
Heatmap of 23 differentially expressed (DE) miRNAs in mature ovaries after exposing zebrafish to high temperature during sex differentiation. The color scale ranges from blue to red, where blue shows relative overexpression and red is relative underexpression. Both miRNAs and samples were grouped by hierarchical clustering.

### miRNA target predictions and functional annotation

To inspect the biological roles of the identified 24 DE miRNAs after high-temperature treatments in the zebrafish gonads, target genes of the 24 miRNAs were predicted upon the zebrafish genome by 3’-UTRs. Most of the DE miRNAs had multiple target genes and many of them were regulated by more than one miRNA. We predicted 1,205 and 101 target genes for ovary and testis, respectively, in the control groups. In ovary, 407 unique targets were found for the 23 DE miRNAs whereas in testis, 85 unique targets were found for dre-miR-122-5p. The full list of the predicted targets is shown in S4 Table.

To better understand the relationship between DE miRNAs and their function in the gonads after heat exposure, GO enrichment analyses of the putative target genes were performed (S5 Table). In ovary, 54 GO terms for Biological process (BP), 27 for Cellular component (CC), and 42 for Molecular function (MF) were predicted and 3 GO terms for BP, 4 for CC, and 3 for MF in testis. In ovary, some of the most enriched GO terms for BP were: regulation of transcription, signal transduction and transport (Fig 3A); for CC were: membrane, nucleus and integral component of membrane (Fig 3B), and for MF were: metal ion binding, zinc-binding, and transferase activity (Fig 3C).

**Fig 3.**
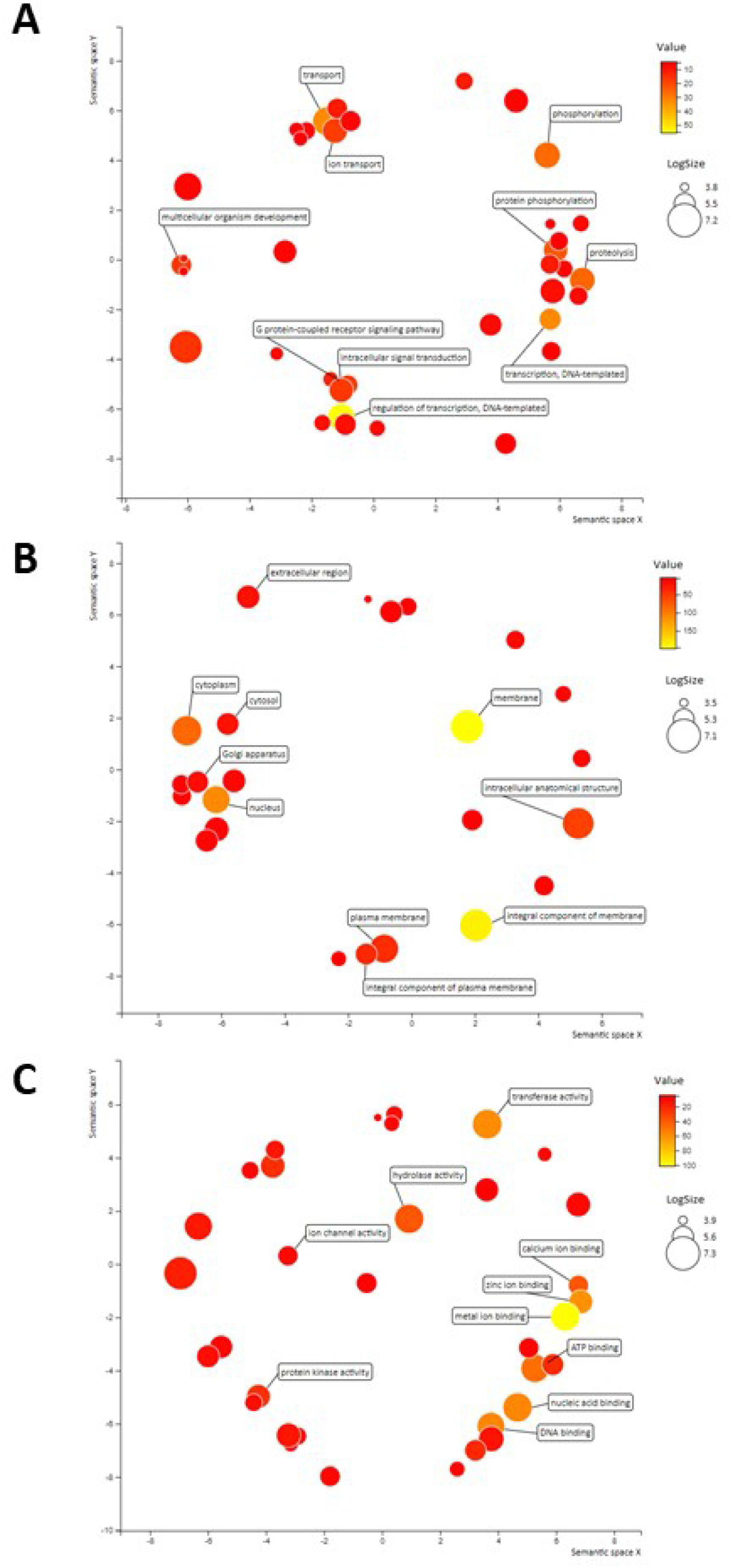
Visual representation of Gene Ontology (GO) terms obtained from predicted target genes of differentially expressed miRNAs in ovary. Color intensity represents the frequency of the GO term as linked to the target genes. LogSize shows the frequency of the GO term in the UniProt database. The top 10 most frequent GO terms are annotated in the plot. **A)** GO terms related to Biological processes. The most frequent terms were regulation of transcription, signal transduction and transport. **B)** GO terms related to Cellular components. The most frequent terms were membrane, nucleus and integral component of membrane. **C)** GO terms related to Molecular function. The most frequent terms were metal ion binding, zinc-binding, and transferase activity.

### Synteny of the target genes

A synteny map indicated the widespread distribution of the 24 DE miRNAs (23 in ovaries, 1 in testis) in the zebrafish genome (Fig 4). In the ovary, the spatial distribution of the 407 target genes in which DE miRNAs interacted was mostly localized in chromosomes 7, 2, 4, 3 and 11 (Fig 5A). These five chromosomes contained 54.5% of the predicted target genes. 16 DE miRNAs are targeting genes in chromosome 4 (S6 Table), some of which were related to the reproductive system (e.g. SRY-box transcription factor 5, *sox5*, RAS like estrogen-regulated growth inhibitor, *rerg*) or the immune system (interleukin 15 receptor subunit alpha, *ilr15β*, B-cell translocation gene 1, *btg1*). In testis, Fig 5B shows the top 15 chromosomes in which the 85 predicted genes were localized. The chromosomes 14, 2, 7, 15, 1 contained 35.3% of the predicted genes.

**Fig 4.**
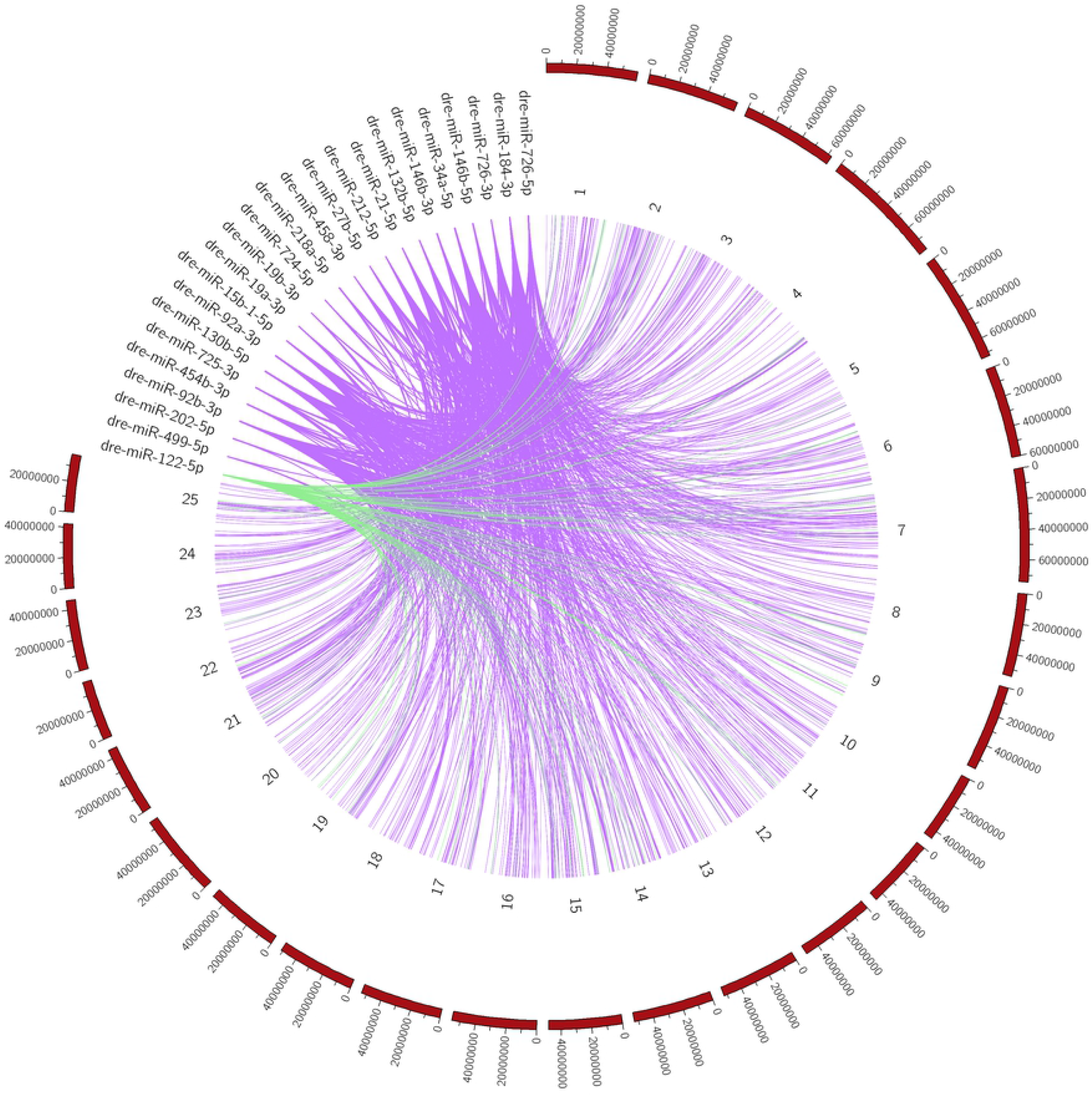
Circular localization of predicted target genes from differentially expressed (DE) miRNAs in the zebrafish genome. 407 predicted target genes of DE miRNAs in the ovary (purple) and 85 predicted target genes in the testis (green) were distributed throughout the zebrafish genome, with the highest percentage present in chromosomes 7 and 14 for ovary and testis, respectively.

**Fig 5.**
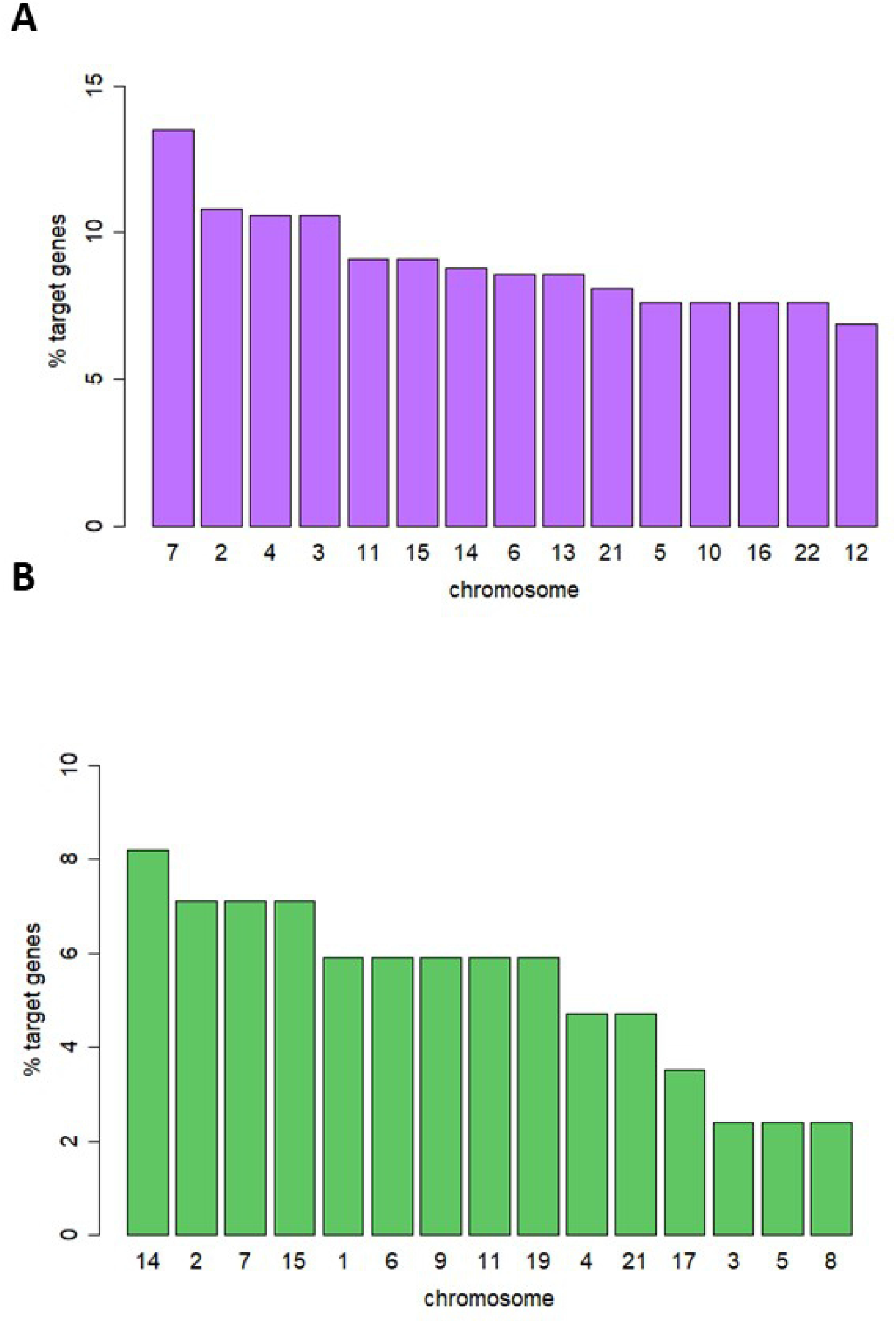
Chromosomal distribution in the zebrafish genome of the number of predicted target genes obtained from differentially expressed miRNAs after high temperature in the ovary (A) and testis (B) in adult zebrafish.

### Spatial expression of selected miRNAs in the gonads

To better understand the functionality of the DE miRNAs in the gonads, the cellular localization of two DE miRNAs was performed by FISH. dre-miR-146b-5p was selected since it was DE in both OHT vs. OCT and OCT vs. TCT in the present data, as well as DE in ovary vs. testis in the data from Presslauer et al. [17]. dre-miR-122-5p was selected for being the only DE miRNA in testes after high temperature. In ovaries, dre-miR-146b-5p and dre-miR-122-5p were detected in the germ cells with an expression that was inversely proportional to oocyte maturation (Fig 6 and 7), detecting expression in small oocytes likely corresponding to pre-vitellogenic and early-vitellogenic oocytes, whereas no expression was detected in large oocytes, including late and post-vitellogenic oocytes. No signal was detected in any of the follicular cells of the ovary. In testis, the signal was not detected for any of the probes used, neither in germ nor follicular cells (data not shown). No signal was detected in negative controls, where scramble miRNA probes were used.

**Fig 6.**
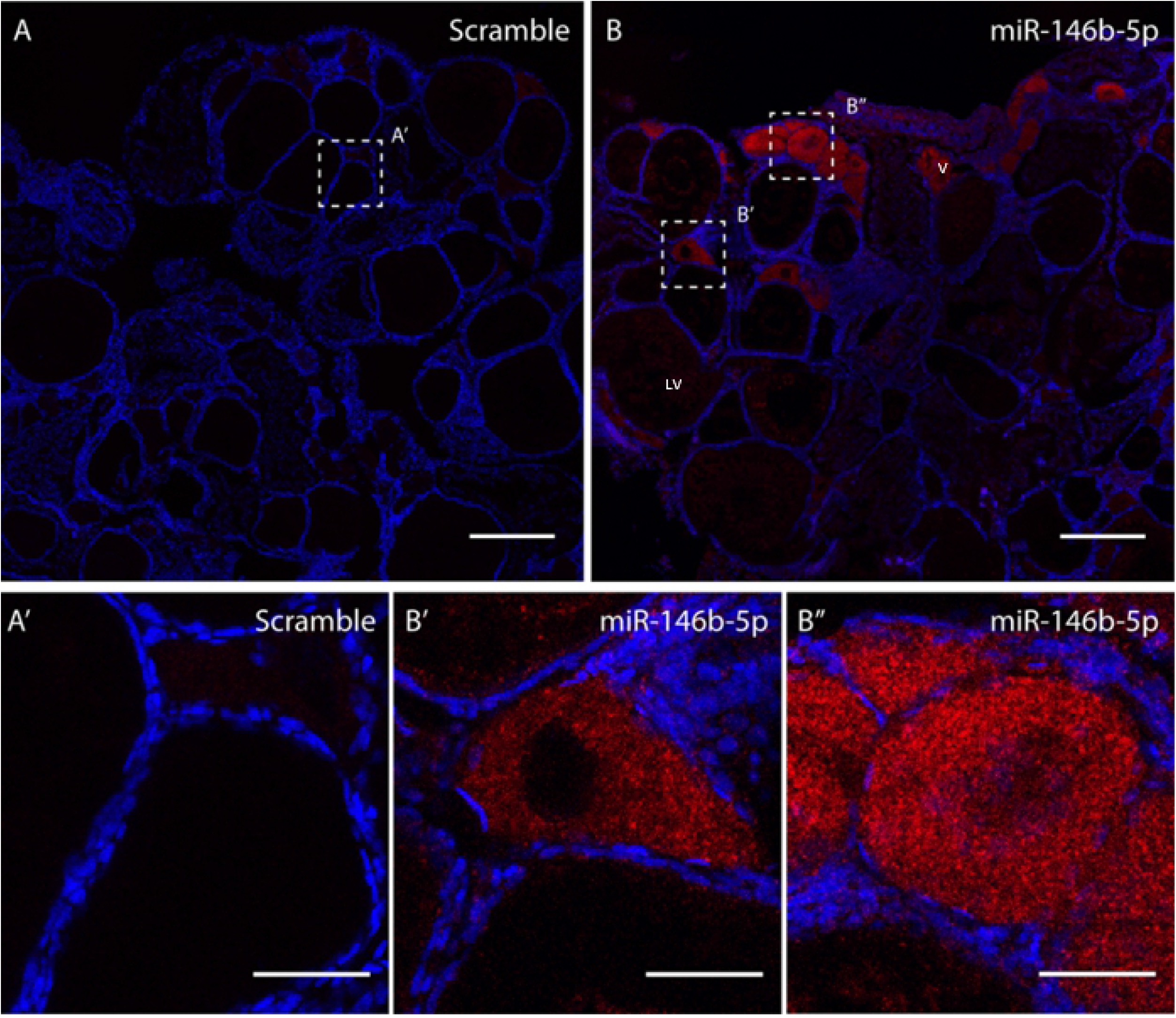
Fluorescent *in situ* hybridization (FISH) of dre-miR-146b-5p in the ovary of adult zebrafish. A total of 6 female fish were used to obtain the results. **A)** Sections **A** and **A’** showed scramble probe. Section **B, B’** and **B”** showed the localization of miR-146b in germ cells. size bar = 100 μm.

**Fig 7.**
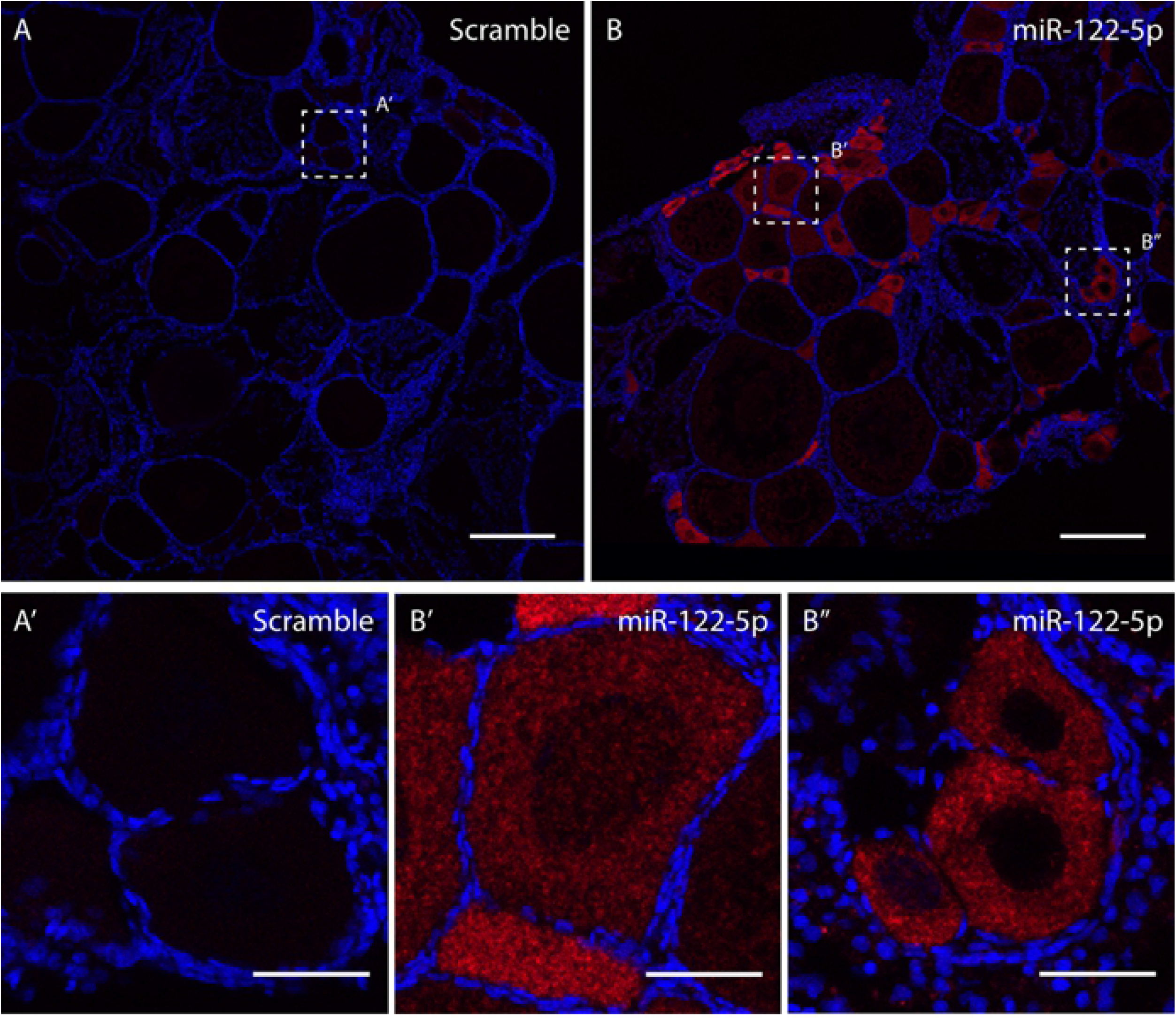
Fluorescent *in situ* hybridization (FISH) of dre-miR-122-5p in the ovary of adult zebrafish. A total of 6 female fish were used to obtain the results. Sections **A** and **A’** showed scramble probe. Section **B, B’** and **B’’** showed the localization of dre-miR-122-5p in germ cells. (V) Vitellogenic oocyte, (LV) late vitelogenic oocyte. A and B scale bar = 500 μm and A’, B’, B” size bar = 100 μm.

## Discussion

Temperature increase influences sexual development by skewing sex ratios towards males in fish [5]. The study of the underlying epigenetic mechanisms of this masculinization have relied on DNA methylation analyses in the gonads of some fish species, like European sea bass [6,37], half-smooth tongue sole [8], tilapia [38], fugu [7] and zebrafish [9]. However, other epigenetic mechanisms, and specifically translation interference by miRNAs, have not yet been elucidated. Here, gonadal data is described on miRNAs affected by changes in temperature during early development that likely play a role in the final sexual phenotype in zebrafish.

To date, available reports in zebrafish show high variability on the sex ratio changes in zebrafish subjected to high temperatures (from 22 to 60% masculinization) [27,39,40], as zebrafish present interfamily variation due to the genetic and environmental influences on the final sexual phenotype [41,42]. In the current study, a 17% sex-reversal was observed (from 70% at low temperature to 87% at high temperature), although not significant. Non-significance might be explained by a low number of biological replicates or/and by the genetic factor of the skewed sex ratios towards males of the family used (i.e. 70%) [43]. To induce a significant masculinization, higher temperatures (i.e. 36°C) can be performed but in contrast, few or no female samples would have been obtained [9,27]. Thus, the experimental approach confirmed the successful implementation of heat treatment as well as validating previously reported results.

Here, we reported a total of 24 DE miRNAs in the zebrafish gonads two months after heat exposure during fifteen days of early development when gonads are differentiating. In a similar study in Atlantic cod, embryos were incubated at high temperatures resulting in alterations of some, but few, miRNAs in juvenile animals in different tissues, including gonads, although the sexual dimorphic difference was not studied [25]. Thus, the alteration of the miRNA expression due to environmental cues indicates that they can be considered as heat recorders as their expression depends on past events. Only one out of the 24 miRNAs altered by elevated temperatures is testis specific. By using the same experimental approach in zebrafish in Ribas *et al*. 2017, testicular transcriptome presented no DE genes after the heat exposition during sex differentiation when compared to the control, revealing that in testes, of some certain neomales, no relevant transcriptomic differences after the heat treatment was presented [27]. Nevertheless, in the same study, another neomale population in the heated group showed a larger amount of DE genes (~700) when compared to the control. In the ovarian transcriptomes, only 20 DE genes were found when compared to the control but a larger number of DE genes were found (~9,650) when compared to so-called *pseudofemales* (females with phenotypic ovaries and with a male-transcriptomic profile). Overall, when comparing the overviewed number of DE miRNAs and the DE genes obtained from both studies in the zebrafish gonads treated with high temperature during sex differentiation, the alterations in the miRNome and the transcriptomes were more severe in the ovaries, probably due to the fact that in the adult female fish, ovaries needed to resist the sex-reversal process while some adult males were already sex-reversed females.

Here, twelve of the miRNAs were upregulated in the adult ovaries of the heat-treated fish, among them, dre-miR-202-5p. Emerging evidence suggests that this miRNA is highly expressed in female gonads of many animals, e.g., fish [25,44], frogs [45] and goats [46]. Although it was proposed as a regulator of fish fecundity and fertility [47], in mammals, miRNA-202 was found in testes in both Sertoli cells and spermatogonia stem cells [48,49] and in rainbow trout more abundantly in testes than ovaries (20 and 10%, respectively) [16] as well as in medaka [47]. Another miRNA that was upregulated was dre-miR-92a, and has been found to be the most abundant miRNA in zebrafish gonads [17]. It was responsible for cell cycle progression during the early stage of embryo development and metamorphosis in Japanese flounder (*Paralichthys olivaceus*) [50] and in zebrafish [51].

The expression of eleven miRNAs was downregulated due to the temperature increase, for example, dre-miRNA-21-5p. This miRNA is highly conserved throughout evolution and abundantly distributed in many tissues in fish. This is the case of the heart [52], kidney [53], and ovary [54]. It was also linked to the fish immune response through the TLR28 signaling pathway [55]. Many functions have been related to miRNA-21 in humans as being found in different cancers, although in fish, fewer data of its biological role are available. Strikingly, most of the miRNAs here identified as heat recorders, are related to ovarian and prostate cancer in humans, either promoting or suppressing cancer progression and thus much literature related to these diseases is available. This is the case of, for example, miR-19b [56], miR-15b [57], and miR-454 [58,59] three upregulated miRNAs in the fish ovaries after the heat and; miR-27b [60,61], miR-212 [62], miR-146b [63], and miR-34a [64], which were downregulated. The emerging research on miRNAs has flourished the utility of miRNAs as bioclinical markers in human cancers during the last decade [65,66] but also as attractive drug targets for human diseases with no current effective treatments [67,68]. This has attracted the attention of many pharmaceutical companies which are developing clinical trials, such as, miRNA-21 and mRNA-92 which are in phase 2 and 1, respectively [68,69] and were found down- and upregulated, respectively, in the ovaries after the heat in the present study. Thus, the exploration of the usefulness of miRNAs as heat markers becomes attractive as a potential method to predict animals with different susceptibilities to environmental cues.

Predicted target genes from the DE miRNAs due to exposure to elevated temperatures, showed functions related to reproduction and sex. This is the case of Polycomb Group RING Finger (*pcgf6*) gene-targeted by dre-miR-458-3p [70];*pcgf5a* gene targeted by dre-miR-184-3p [71], and Dickkopf-related protein (*dkk1b*) targeted by dre-miR-212-5p [72]. Similarly, those miRNAs downregulated in the ovaries after the heat targeted to reproduction-related genes such as *sox5* targeted by dre-miR-15b-1-5p [73], and Nuclear Receptor Subfamily 5 Group A Member 2 (*nr5a2*) targeted by dre-miR-19b-3p [74]. In the testes, only one miRNA was identified as heat recorder, miR-146b and targeted, for example, to *bbc3* gene which is related to prostate cancer [75], and *tet2*, a demethylator of many genes, included the SRY, a key gene in the regulation of male sex determination in mammals [76]. Overall results confirmed that the miRNA machinery was active and essential to regulate the environmental cues that occur in the adult fish gonads.

The synteny of the predicted target genes of the heat recorders miRNAs on the zebrafish genome showed multiple regions in all the 25 chromosome pairs, but more abundantly in chromosomes 7, 2, 4, 3 and 11 in the ovary, accounting for 54.5% of the predicted target genes in the present data. To foster the identification of sex-determining gene(s) in this popular animal model, many sex genetic studies in the last decade have been performed by crossing natural and domesticated zebrafish strains. By single nucleotide polymorphisms (SNPs) and sequence-based polymorphic restriction site associated (RAD-tag) strategies, several sex-linked loci in the chromosomes 4 and 3 have been identified [77,78] and in the chromosomes 5 and 16 [78,79]. Strikingly, in chromosome 4, the sex-association region (*sar*) was localized in wild zebrafish strains [80], a chromosome that from our data supported more than 10,5% of the predicted target genes of the miRNAs sensitive to heat. Furthermore, chromosome 16 accounted ~7,5% of our predicted target genes. Thus, although more research is required to understand the biological functions of the present data, we can ascertain some of the chromosomes that host genes regulated by miRNAs sensitive to heat.

Further, we identified sexual dimorphism in the expression of miRNAs in the fish gonads with a total of 45 and 54 up- and downregulated, respectively, in the ovary when compared to testis. To increase consistency, our data were compared with two available data of the same species resulting in common miRNAs. We found that miRNA-200b-3p was upregulated in the ovary in the three zebrafish comparisons. The role of this miRNA is not fully understood but it is known to be involved in many human cancers: kidney [81], prostate [82], and breast [83]. It is highly released in the serum of the anovulatory women diagnosed with polycystic ovary syndrome and suggested as a clinical marker [84]. miR-212-5p was DE in the testes *vs* ovaries in the three zebrafish gonadal miRNA datasets and inhibited after the heat treatment in the ovaries. This can indicate its role by dysregulation of ovarian functions during the masculinization event occurred by heat. The miRNA-212 function is not stated but in humans, it is related to cell proliferation and angiogenesis and is present in the brain and gonads [62,85]. In addition, miRNA-212 was found in tilapia gonads [86], which, together with the current results, show its relevance presence in the reproduction system in fish.

The gonadal localization of two miRNAs, dre-miR-122-5p and dre-miR-146b-5p, showed similar results. In ovary, their expressions were found in the germ cells but not in the granulosa or theca cells while fluorescent intensity was stronger in less mature oogonial cells, suggesting a potential role of these miRNAs in germ cell development. In testis, although miR-122 is involved in zebrafish sperm quality [87,88] and male fertility in mammals [89], its localization, together with that for miR-146b, was not possible in the zebrafish testicular cells, so more sensitive methods need to be readied. miR-122 is involved in humans in many cancer and has reached phase II in clinical trials for treating hepatitis [67,90]. In fish, much literature related to miR-122 is available certifying the role in the immune [91,92] and in the metabolic systems being highly abundant in the zebrafish liver [93]. The presence of miR-122 was detailed in many fish species such as tilapia, medaka, carp, and in many fish tissues such as the spleen, head kidney [94–97]. In the gonads, it was detected in mature sharpsnout seabream (*Diplodus puntazzo*) but not in the marine medaka [95,98]. Strikingly, miRNA122 was reported to be sensitive to cold temperatures in the Senegalese sole (*Solea senegalensis*) [24,99], thus the role of this miRNA as a thermal recorder is worth further exploring. Regarding miR-146b in humans, it plays a role in the innate immune response [100] and is involved in gliomas and ovarian cancers [101,102]. In fish, very little data is available but it was upregulated in response to infection in zebrafish embryos [103] and spleen [104] and the sperm of growth hormone (GH)-transgenic zebrafish [87]. Overall, to our knowledge, this is the first time that the cellular localization of these two miRNAs are described in the gonads.

## Conclusions

Present data evidence that high temperature alters the miRNome in the fish gonads. The influence of heat treatment during gonadal development altered the expression of 23 miRNAs in the ovaries, by enhancing, for example, miR-92b-3p and miR-202-5p, or repressing, for example, miR-212-5p and miR-146b-3p expressions. In testes, miR-122-5p was the only miRNA sensitive to heat. These miRNAs act as heat recorders and might be potential targets for developing predictive tools of heat response, essential in a climate change scenario or to increase productivity from sustainability. In addition, as most of the 24 DE miRNA have been found to be involved in different diseases, but mostly related to cancer, the data here might be helpful to enhance our knowledge on the functional roles of the miRNAs identified in the present study.

## Acknowledgments

This study was supported by the Spanish Ministry of Science grant AGL2015-73864-JIN “Ambisex” and 2PID2020-113781RB-I00 “MicroMet” and by the Consejo Superior de Investigaciones Científicas (CSIC) grant 02030E004 “Interomics” to LR, by grant AGL2016-78710-R to FP and with funding from the Spanish government through the ‘Severo Ochoa Centre of Excellence’ accreditation (CEX2019-000928-S). We thank the lab technician Sílvia Joly for her essential assistance in our team, Alejandro Valdivieso for the treatments of the zebrafish, the master student Irene Santisteban for helping with some miRNA analyses and Gemma Fusté for her assistance in fish facilities.

## Ethics

Experimental procedures agreed with the European regulations of animal welfare (ETS N8 123,01/01/91) and obtained approval with project number 9977 by the Catalan government regulations (34, 53/2013).

## Supplementary tables

S1 Table. Weight, length and K-factor of all zebrafish used for the experiment.

S2 Table. Common miRNAs in ovaries and testes between present data and Presslauer *et al* 2017 and Desvignes *et al* 2019.

S3 Table. Statistical data of 24 differentially expressed miRNAs in the zebrafish mature gonads after high-temperature treatment during sex differentiation.

S4 Table. Predicted target genes of the 24 differentially expressed miRNAs

S5 Table. Gene Ontology (GO) terms obtained from the predicted target genes.

S6 Table. Differentially expressed miRNA in the ovaries after high temperatures found in chromosome 4.

## Supplementary figures

**S1 Fig. Sex ratio in adult zebrafish after high temperature (34°C) treatment during sex differentiation.** Results show the mean ± SD of two technical replicates of one family pair for control (CT, 28°C; n = 27) and treated (HT, 34°C; n = 52) groups. No significant differences were found between the two groups by Chi-squared test.

**S2 Fig. Multidimensional scaling of ovary and testis RNA sequencing data from 16 samples.**Four in each group, ovary and testis, control and high temperature.

**S3 Fig. Validation of RNA sequencing data by qPCR.** The comparison is based on the log2 fold. The miRNAs compared were for OHT *vs*. OCT: dre-miR-202-5p, dre-miR-92a-3p, dre-miR-21-5p and dre-miR-146b-3p; for THT *vs*. TCT: dre-miR-122-5p and for OCT *vs*. TCT: dre-miR-146b-3p and dre-miR-2189-3p.

**S4 Fig. Venn diagrams of differentially expressed (DE) miRNAs in ovary *vs*. testis. A)** Common DE expressed miRNAs in ovaries between Presslauer *et al* 2019, Desvignes *et al* 2017, and present data. One miRNA was DE in all datasets: dre-miR-200b-30. **B)** Common DE expressed miRNAs in testes between Presslauer *et al* 2019, Desvignes *et al* 2017, and present data. One miRNA was DE in all datasets: dre-miR-212-5p.

**S5. Fig Top five up- and downregulated miRNAs in adult ovaries heated with high temperature in zebrafish.**

## Datasets

**Dataset 1**. Reads of aligned sequences obtained in the ovaries and testes in adult zebrafish control group and treated with high temperature.

